# Social reward processing and financial vulnerability in older adults vary with environmental stressors and social support

**DOI:** 10.64898/2026.11.27.769758

**Authors:** Katherine Rodman, Jacob M. Stanley, Sophia DiConza, Ryan S. Gephart, Cooper J. Sharp, Jeremy Mennis, Ingrid R. Olson, Dominic S. Fareri, Tania Giovannetti, David V. Smith

## Abstract

Social rewards play an important role in motivating behavior and guiding decisions in older adulthood. Changes in the neural systems supporting social reward processing may have consequences for financial decision making, including vulnerability to exploitation. Although environmental stressors have been implicated in cognitive decline, little is known about their associations with social reward processing or whether social resources moderate these associations. We examined these questions in 97 adults aged 55–89 years who completed matched social and monetary reward tasks during functional magnetic resonance imaging. Perceived social support moderated the association between neighborhood deprivation and neural responses to social relative to monetary rewards near the right temporoparietal junction. Greater deprivation was associated with smaller responses to social relative to monetary wins at lower levels of perceived support, whereas this association reversed at higher levels of support. Perceived support also moderated the association between long-term exposure to fine particulate matter and financial exploitation vulnerability, with greater exposure associated with greater vulnerability among older adults reporting less support. Moreover, financial exploitation vulnerability was associated with differences in ventral striatal connectivity during social relative to monetary feedback, although this association was sensitive to adjustment for perceived support. Together, these findings extend research on environmental correlates of brain health to neural systems supporting social reward processing. Understanding individual differences in these systems and their relevance to financial vulnerability may require considering the environmental conditions in which older adults live alongside the social resources available to them.

**Significance:** Social rewards motivate behavior throughout adulthood and may be particularly important in later life, when financial decisions can have lasting consequences for independence and cognitive health. Although environmental stressors have been linked to cognitive decline, their relevance to the neural processing of social rewards remains poorly understood. We show that perceived social support moderates associations between neighborhood deprivation and neural responses to social relative to monetary rewards in older adults. Perceived support similarly moderates the association between air pollution and financial exploitation vulnerability. These findings highlight the importance of considering environmental and psychosocial conditions when investigating the neural basis of social and economic decision making in older adulthood.

## 1. Introduction

Financial exploitation threatens older adults’ financial independence and well-being. Approximately 1 in 18 older adults experience financial fraud or scams annually (Burnes et al., 2017). Age-related changes in executive function, episodic memory, and social cognition may increase vulnerability to exploitation (Spreng et al., 2016; Fenton et al., 2022). Indeed, lower scam awareness has been associated with subsequent cognitive impairment, dementia, and greater Alzheimer’s disease pathology (Boyle et al., 2019). Substantial losses of wealth have also been associated with cognitive decline and elevated dementia risk, although cognitive changes may precede financial losses (Pan et al., 2023; Cho et al., 2026). Understanding the neural basis of financial exploitation vulnerability may therefore be important for healthy aging.

Among the neural systems relevant to financial exploitation vulnerability are those supporting social reward processing. Socioemotional selectivity theory posits that people prioritize emotionally meaningful relationships as their perceived time horizons shorten (Carstensen et al., 1999; Carstensen, 2006). Consistent with this account, older adults exhibit greater preferences for immediate social rewards (Seaman et al., 2016) and heightened striatal responses during anticipation of social relative to monetary rewards (Rademacher et al., 2014). However, relative to younger adults, older adults exhibit less striatal sensitivity to partner closeness during reciprocated trust and different patterns of connectivity between the default mode network and temporoparietal junction (Fareri et al., 2022). Individual differences in social reward processing could contribute to financial exploitation vulnerability in several ways. Heightened responsiveness to social rewards may encourage individuals to seek interpersonal experiences or prioritize social approval, increasing opportunities for exploitation. Conversely, reduced differentiation of positive and negative social feedback may limit the ability to use interpersonal information to guide decisions. Indeed, older adults exhibit attenuated anterior insula responses to cues of untrustworthiness (Castle et al., 2012), and financially exploited older adults exhibit structural and functional differences in regions implicated in socioemotional processing (Spreng et al., 2017).

Environmental conditions may contribute to individual differences in these neural systems. Neighborhood deprivation has been associated with poorer cognitive functioning (Wu et al., 2015), accelerated cortical thinning, and cognitive decline (Hunt et al., 2021). Exposure to air pollution, including fine particulate matter (PM2.5), has also been associated with poorer cognitive performance (Chen, 2018; Zhang et al., 2018). Emerging research links environmental disadvantage to differences in neural reward processing earlier in life (reviewed in Polemiti et al., 2024). Whether neighborhood deprivation and air pollution relate to neural responses to social rewards in older adults, and whether these associations vary with everyday cognitive difficulties, remains unclear.

Associations between environmental stressors and brain function may also depend on the social resources available to individuals. Greater social support has been associated with better cognitive functioning in older adulthood (Mogic et al., 2023). Associations between air pollution and cognitive outcomes also vary with social engagement and neighborhood social cohesion (Yuan et al., 2024; Frndak et al., 2024). Although these constructs differ from perceived interpersonal support, these findings suggest that social resources may moderate environmental associations with brain health. Perceived support may be particularly relevant to social reward processing because existing social resources may shape the motivational significance of social acceptance and rejection.

Here, we examined how neighborhood deprivation and air pollution relate to social reward processing and financial exploitation vulnerability in older adults. Social and monetary rewards engage overlapping neural circuitry (Izuma et al., 2008). By comparing social with monetary wins and comparing win–loss responses across domains, we assessed relative responsiveness to social rewards and social modulation of feedback valence, respectively. Using matched reward tasks during functional MRI, we asked three questions. First, do associations between environmental stressors and financial exploitation vulnerability vary with perceived social support or everyday cognitive difficulties? Second, is financial exploitation vulnerability associated with neural social reward responses, including regional activation and ventral striatal connectivity? Third, are environmental stressors associated with social reward processing, and do these associations vary with perceived support or everyday cognitive difficulties?

## 2. Methods

### 2.1. Participants

We drew our sample from an ongoing study of social decision making that recruited participants in the greater Philadelphia area through flyers and online advertisements (Smith et al., 2024). The protocol included a behavioral visit and an fMRI visit. Participants provided written informed consent under a protocol approved by the Temple University Institutional Review Board and were compensated for their time. The baseline imaging dataset included 151 adults aged 55 years or older. Nine had no documented baseline Social Doors acquisition, two had partial runs, and one made no recorded choices. Of the 139 with usable task data, two did not meet the 95% reference-mask coverage criterion, leaving 137 participants. Two had no matching screener record, 24 had incomplete demographic, questionnaire, or address data, and one lacked a numeric Area Deprivation Index rank. All 110 remaining participants had complete residential PM2.5 data. We required both task runs to pass quality control: 11 participants exceeded the mean framewise displacement threshold in at least one run (1.5 interquartile ranges above the third quartile for that task and session), and two had less than 95% reference-mask coverage on the monetary run. Low temporal signal-to-noise ratio accounted for no additional exclusions. The final analytic sample included 97 participants.

### 2.2. Experimental procedure

During the initial visit, we photographed participants and told them that age-matched peers would review their photograph and indicate whether they liked or disliked them. This cover story helped convince participants they would receive real feedback at the scan visit; the photographs were not shared with peers and were deleted. Several days later, participants performed the social feedback task during fMRI and received feedback they believed was from these peers.

### 2.3. Social and monetary feedback tasks

Participants completed two matched guessing tasks: Social Doors, which delivered purported peer-acceptance or peer-rejection feedback, and Doors, which delivered monetary win or loss feedback (Distefano et al., 2018; Quarmley et al., 2019; Smith et al., 2024). In Social Doors, participants viewed two peer faces and selected the person they believed had liked them. A green check mark indicated positive feedback (the selected peer supposedly liked them), whereas a red X indicated negative feedback. In Doors, participants selected between two doors and received similar feedback indicating a monetary win or loss. Each participant completed one run of each task, with run order counterbalanced.

Each run contained 60 trials, divided equally between positive and negative feedback. A trial included a 3 second decision display and 1 second feedback display, separated from the next trial by a variable intertrial interval (1.1–11.6 s; mean, 3.5 s). Each run comprised 292 functional volumes (Figure 1). The two tasks therefore matched the frequency and timing of feedback valence while varying whether feedback concerned social acceptance or a monetary outcome.

**Figure 1.**
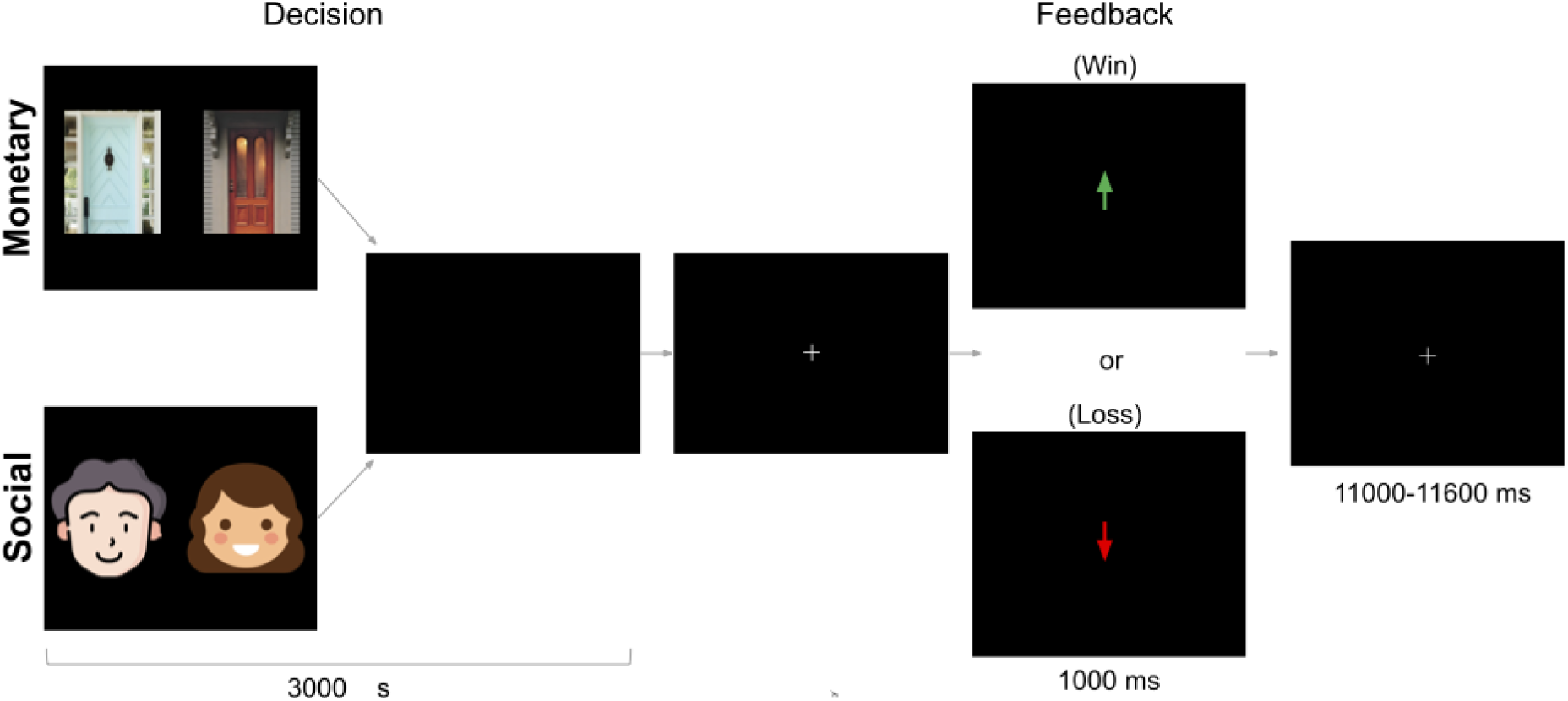
Schematic of the Social Doors (social reward) and Doors (nonsocial reward) tasks. During the decision phase, participants selected between two stimuli (faces for the social condition and doors for the nonsocial condition). Pictures of real faces were used in the experiment. Following a fixation period, feedback was presented indicating a win or loss. Neural activity was measured during the feedback phase. Figure adapted from Smith et al. (2024). Face icons were obtained from Flaticon (Sudowoodo, n.d.; Magnific, n.d.).

### 2.4. Neighborhood and individual-difference measures

We indexed neighborhood socioeconomic disadvantage using the 2024 national Area Deprivation Index (ADI), linked to participants’ residential locations; higher scores represent greater area-level deprivation (Kind C Buckingham, 2018). Air pollution was measured using mean residential fine particulate matter (PM2.5) concentrations across the 2013-2023 period. We assessed perceived social support using the corrected score from the Multidimensional Scale of Perceived Social Support (MSPSS; Zimet et al., 1988), with higher scores indicating more perceived support. The Everyday Cognition scale (ECog; Farias et al., 2008) assessed self-reported change in everyday cognitive functioning; higher scores indicate greater perceived decline rather than a clinical diagnosis. We used the Financial Exploitation Vulnerability Scale (FEVS; Hall et al., 2022) to examine a self-reported measure of risk for financial exploitation. Higher scores indicate a greater risk for financial vulnerability.

### 2.5. Behavioral analyses

We used linear regression to examine whether perceived social support and everyday cognitive difficulties moderated associations between environmental stressors and financial exploitation vulnerability. In separate models, we regressed financial exploitation vulnerability scores on ADI or PM2.5, the corresponding moderator, and their interaction, adjusting for age and gender. Continuous predictors were mean-centered before constructing interaction terms. We estimated confidence intervals and conducted inference using heteroskedasticity-consistent (HC3) standard errors. We adjusted for multiple comparisons using the Holm procedure within families of related tests. To characterize interactions, we estimated conditional associations between environmental exposures and financial exploitation vulnerability across observed levels of the moderators. Additional models examined whether these associations persisted after adjustment for the other environmental exposure and individual-difference measures.

### 2.6. MRI acquisition and preprocessing

We acquired images at the Temple University Brain Research and Imaging Center using a 3 T Siemens Prisma scanner and 20-channel head coil (Smith et al., 2024; Wang et al., 2026). The four-echo functional echo-planar imaging sequence used echo times of 13.8, 31.54, 49.28, and 67.02 ms, a repetition time of 1,615 ms, multiband factor of 3, in-plane GRAPPA acceleration of 2, 51 axial slices, and 2.7-mm isotropic voxels with a 10% interslice gap. We also acquired single-band reference images for registration and high-resolution T1-weighted anatomical images (MPRAGE; 1-mm isotropic resolution).

We converted the images to Brain Imaging Data Structure (BIDS) format using HeuDiConv (Halchenko et al., 2024) and preprocessed the data using fMRIPrep 25.2.5 (Esteban et al., 2019). We generated phase-derived field maps with MEDIC to correct susceptibility-related distortion and used tedana for multi-echo combination and denoising (DuPre et al., 2021; Van et al., 2026). We analyzed functional data in MNI152NLin6Asym space and applied 5-mm full-width-at-half-maximum Gaussian smoothing in FSL. Detailed anatomical and functional preprocessing, multi-echo processing, distortion correction, and quality-control procedures appear in the Supplementary Methods.

### 2.7. Neuroimaging analyses

We used FSL FEAT to estimate responses to positive and negative feedback separately in each task (Jenkinson et al., 2012). For each run, we modeled positive and negative feedback events using their recorded onset times and durations, then convolved the event regressors with FSL’s double-gamma hemodynamic response function. The run-level positive-minus-negative contrast measured sensitivity to feedback valence within each domain. Our primary outcome isolated social-relative feedback-valence sensitivity by subtracting the corresponding monetary contrast: [(Social Positive − Social Negative) − (Monetary Positive − Monetary Negative)]. A positive value indicates greater positive-versus-negative differentiation for social than monetary feedback; it does not necessarily indicate greater overall activation during the social task.

We included six rigid-body motion parameters, the first six anatomical CompCor components, non-steady-state volume regressors, framewise displacement, and discrete cosine basis functions generated by fMRIPrep as nuisance regressors. These cosine terms implemented high-pass filtering with a 128-s cutoff, without additional temporal filtering in FEAT. We used FILM prewhitening to account for temporal autocorrelation.

We examined task-dependent connectivity using generalized psychophysiological interaction (gPPI) models with the ventral striatum as the seed (McLaren et al., 2012). For each task run, we combined the feedback-event regressors with the ventral striatal time series and their interaction terms. We contrasted connectivity associated with positive versus negative feedback within each task, then compared the social and monetary contrasts to quantify social-relative ventral striatal connectivity. These contrasts paralleled our activation analyses.

For each participant, we subtracted the monetary COPE from the corresponding social COPE to compare the win–loss, winning, and losing responses; we also averaged the win–loss contrasts across tasks. For activation and ventral-striatum-seeded connectivity, we fit exploratory group-level regression models in FSL randomise. Separate models examined ADI and PM2.5, their interactions with perceived social support or everyday cognitive difficulties, and associations with financial exploitation vulnerability, including sensitivity models. We included both constituent main effects in interaction models, centered continuous predictors, and adjusted for age, gender, scanner-software indicators, and signed between-task FD and tSNR differences (pair means for averaged outcomes). We used 10,000 permutations and, excepted where noted, a cluster-forming threshold of Z > 3.1, testing positive and negative contrasts separately and controlling cluster-level familywise error at p < .05 within each map. Neuroimaging results are shown using MRIcroGL (Rorden, 2025).

## 3. Results

### 3.1 Environmental stressors and financial vulnerability

We asked whether perceived social support and self-reported everyday cognitive difficulties distinguish how environmental stressors relate to vulnerability to financial exploitation. In 97 adults aged 55–89, we compared the associations of fine particulate matter (PM₂.₅) and neighborhood deprivation with scores on the Financial Exploitation Vulnerability Scale (FEVS) across levels of support and everyday cognitive difficulties. Higher scores indicate greater vulnerability on this self-report measure. Greater PM₂.₅ exposure was associated with higher vulnerability at lower levels of support, and the association weakened as support increased (interaction *b* = −0.989, 95% CI [−1.505, −0.473], Holm-adjusted *p* = .00051, Figure 2).

**Figure 2.**
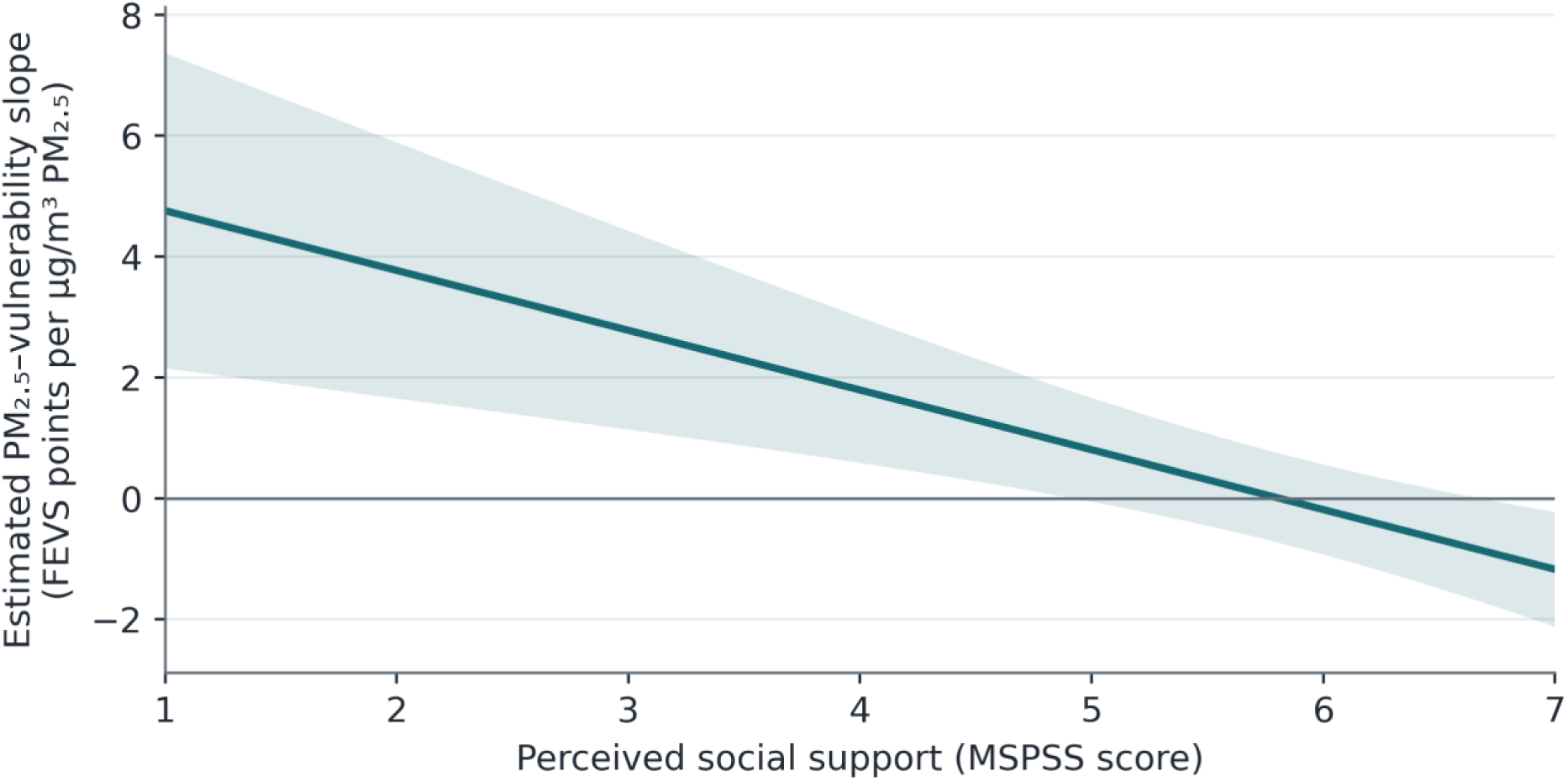
The association between fine particulate matter and financial vulnerability weakens with perceived support. The curve estimates the change in Financial Exploitation Vulnerability Scale score associated with a 1 µg/m³ difference in PM2.5 at each observed level of perceived support. Support was measured with the Multidimensional Scale of Perceived Social Support. The horizontal line marks a slope of zero. Shading denotes pointwise 95% confidence intervals using HC3 standard errors. Estimates are from the age– and gender-adjusted model in 97 adults. Table S2 reports the estimates at representative support values.

Everyday cognitive difficulties could account for some of the variation associated with support. The PM₂.₅-by-support interaction persisted after adjustment for these difficulties (Table S2). There was little evidence that support modified the association with neighborhood deprivation or that everyday cognitive difficulties modified the association with either stressor (Table S1).

### 3.2 Financial vulnerability and neural responses to feedback

Financial vulnerability may relate to how older adults respond to different kinds of feedback. In Social Doors, participants chose which of two peers they thought had liked them and received apparent acceptance or rejection. In the matched monetary task, they chose a door and received a monetary gain or loss. Comparing social wins with monetary wins tests how responses to positive feedback differ across tasks. Comparing the win-minus-loss response between tasks asks whether sensitivity to feedback valence differs across tasks. Wins elicited greater striatal activation than losses in both tasks. The group-average win-minus-loss contrast comparison yielded no corrected clusters.

Because PM2.5 was associated with vulnerability at lower support, we also accounted for exposure in our neuroimaging analyses. We estimated the association with vulnerability at the sample’s mean PM2.5 exposure in models that included an interaction between vulnerability and exposure. We used generalized psychophysiological interaction (gPPI) analysis to estimate how coupling between the ventral striatum and other regions varied with feedback. Higher vulnerability scores were associated with a smaller win-minus-loss difference in ventral striatal coupling with the left frontal pole during social relative to monetary feedback (Figure 3; cluster-corrected *p* = .0303). This association did not survive whole-brain correction after adjustment for perceived social support, although the estimate within the originally identified region changed little (Table S3).

**Figure 3.**
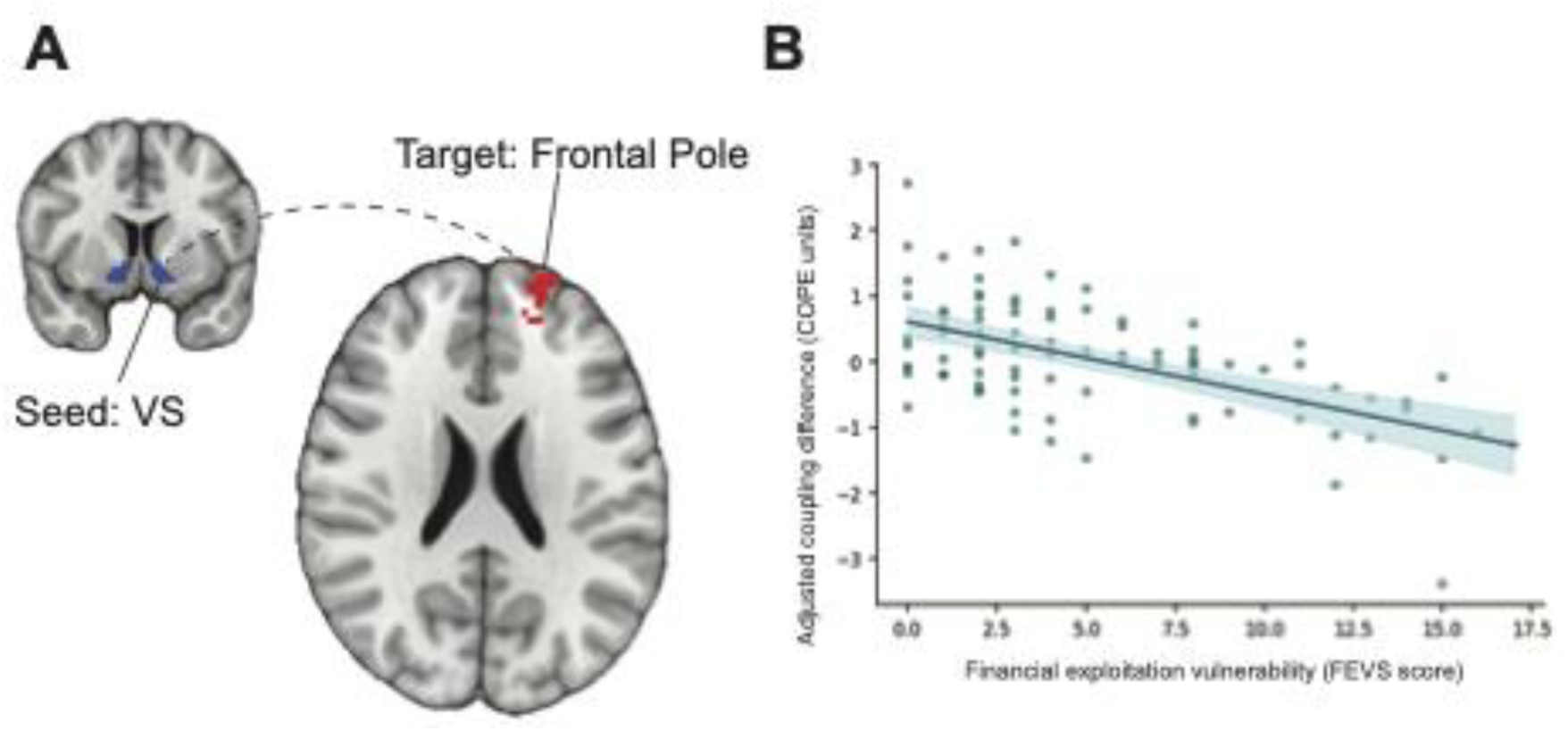
Financial exploitation vulnerability and social modulation of valence-related ventral striatal connectivity. (A) At mean PM2.5 exposure, greater financial exploitation vulnerability was associated with a smaller win–loss difference in connectivity between the ventral striatum and a region near the left frontal pole during social relative to monetary feedback (cluster-corrected p = .0303). (B) Financial exploitation vulnerability scores and covariate-adjusted estimates of the connectivity contrast extracted from the identified cluster. Each point represents one participant.

At mean PM2.5, higher vulnerability scores were associated with greater social-minus-monetary activation during wins in left anterior frontal tissue, much of which the atlas labeled as white matter. The association did not meet the corrected threshold after adding everyday cognitive difficulties alone (Table S4). Including perceived support as a moderator yielded no corrected vulnerability-by-support clusters for activation or ventral striatal coupling.

### 3.3 Environmental stressors and neural responses to feedback

We also asked whether either environmental stressor was associated with neural responses across tasks or with the difference between tasks. Higher Area Deprivation Index (ADI) scores, which reflect greater neighborhood deprivation, were associated with a smaller win-minus-loss difference in ventral striatal coupling averaged across the tasks. The direct associations with deprivation yielded no corrected clusters for activation or for coupling contrasts comparing the tasks. Direct PM2.5 associations yielded no corrected clusters for either neural measure (Table S5).

We next examined whether perceived social support moderated associations between environmental stressors and neural responses to social rewards. The interaction between neighborhood deprivation and perceived support revealed a cluster in right posterior temporal cortex for the contrast comparing social and monetary wins (Figure 4; cluster-corrected *p* = .0255). Greater deprivation was associated with smaller responses to social relative to monetary wins at lower levels of perceived support, whereas this association reversed at higher levels of support. The association with deprivation became more positive with increasing support for social wins and more negative for monetary wins (Table S5, Figure S3). The interaction between deprivation and support did not yield significant clusters for the contrast comparing win– loss responses across tasks. Similarly, interactions between PM2.5 and perceived support did not yield significant clusters for activation or connectivity.

**Figure 4.**
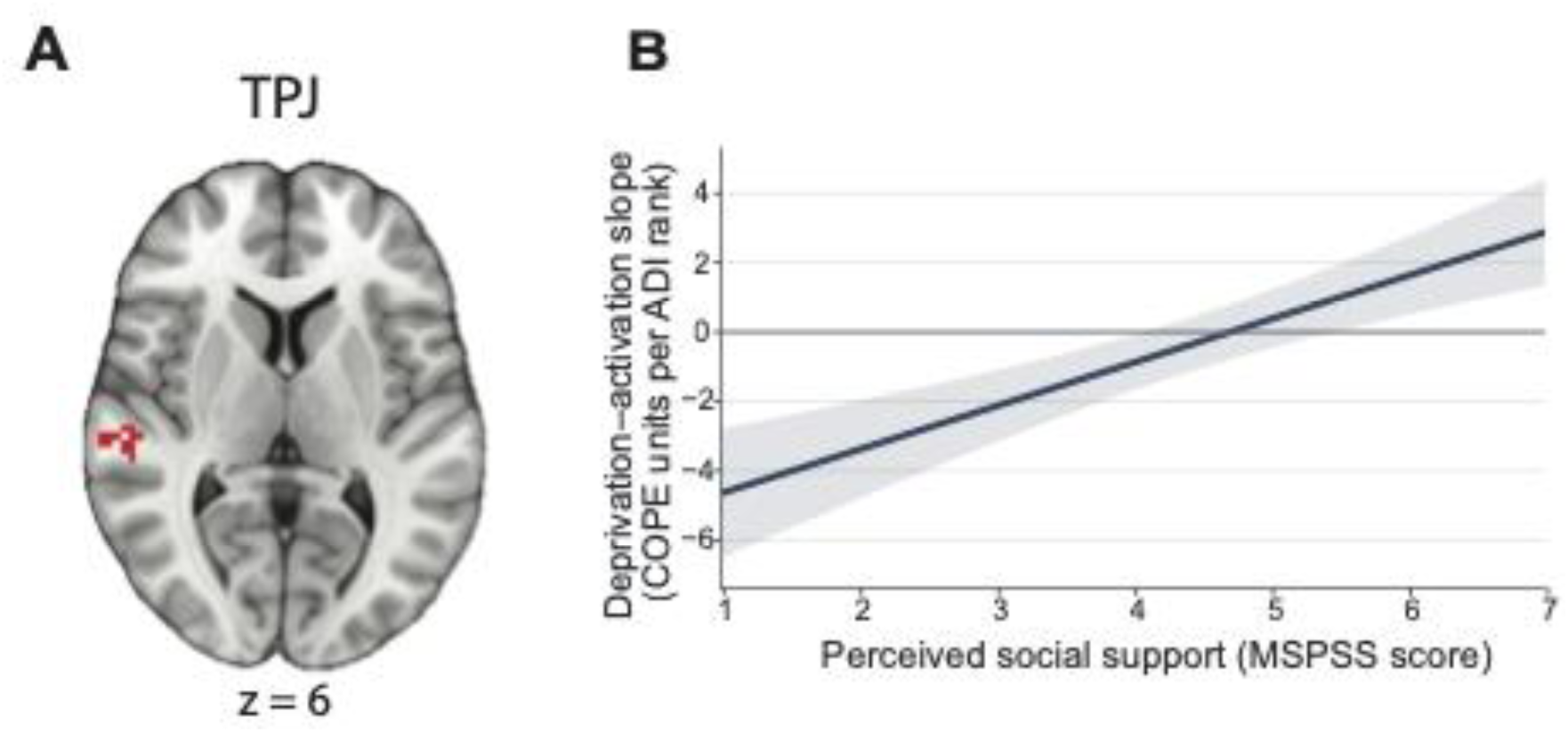
The association of neighborhood deprivation with social relative to monetary winning activation varies with perceived support. (A) The interaction was detected in a right posterior temporal cluster near the temporoparietal junction (cluster-level familywise error *p* = .0255 within the map, cluster-defining threshold equivalent to *Z* = 3.1). The color scale shows the signed voxelwise test statistic. (B) The estimated change in social-minus-monetary winning activation per one-rank increase in Area Deprivation Index, expressed in contrast of parameter estimates (COPE) units. At lower support, higher deprivation was associated with a smaller response to social relative to monetary wins. The slope reversed at higher support. Shading shows pointwise 95% confidence intervals using HC3 standard errors. Estimates illustrate the interaction in the same sample that identified the cluster (*N* = 97).

Everyday cognitive difficulties might also distinguish neural responses at similar exposure levels. The exposure-by-cognition interactions that met correction were confined to coupling. For deprivation, interaction estimates were positive for the contrast comparing the tasks’ win-minus-loss responses and negative for the contrast comparing their responses to losses. The two contrasts showed limited anatomical overlap. For PM2.5, interaction estimates had opposite signs across regions (Tables S6 and S7).

## 4. Discussion

We examined how environmental conditions and perceived social support relate to individual differences in social reward processing and financial exploitation vulnerability in older adults. Greater neighborhood deprivation was associated with smaller neural responses to social relative to monetary rewards in right posterior temporal cortex at lower levels of perceived support and larger relative responses at higher levels of support. Greater PM2.5 exposure was associated with greater financial exploitation vulnerability at lower levels of support, and this association weakened as support increased. Financial exploitation vulnerability was also associated with reduced social modulation of valence-related connectivity between the ventral striatum and a region near the left frontal pole. Together, these findings highlight the importance of considering environmental and psychosocial conditions when investigating the neural basis of financially consequential social decisions in older adulthood.

Our findings complement evidence linking neighborhood disadvantage and air pollution to poorer cognitive outcomes in older adulthood (Wu et al., 2015; Hunt et al., 2021; Chen, 2018; Zhang et al., 2018). Although the earlier studies focused on cognitive health, our results suggest that environmental associations also extend to self-reported financial exploitation vulnerability. The dependence of the PM2.5–vulnerability association on perceived support is consistent with evidence that social support or related social characteristics distinguish cognitive outcomes among adults facing environmental adversity (Mogic et al., 2023; Yuan et al., 2024; Liu C Hu, 2023; Frndak et al., 2024). These studies assess different aspects of social functioning and different outcomes, but together they motivate considering the social circumstances of older adults when evaluating environmental correlates of their well-being. The present findings suggest that the social resources older adults perceive may matter for associations between environmental stressors and outcomes beyond conventional measures of cognitive functioning. Greater air pollution exposure was associated with higher financial exploitation vulnerability at lower levels of support, whereas this association weakened as support increased. Neighborhood deprivation showed a different pattern, varying with perceived support in its association with neural responses to social relative to monetary wins near the right TPJ. The interaction reflected opposing contributions from the two winning conditions, rather than a change confined to social feedback. Thus, comparing social and monetary outcomes revealed contextual differences that would have been difficult to interpret using either task alone. Our findings motivate investigating perceived social support as a source of variability in environmental associations with both financial vulnerability and social-information processing.

Several features of our study constrain the scope of these findings. Residential exposure estimates cannot establish individual exposure histories, and our cross-sectional design cannot determine whether environmental conditions or social resources precede changes in feedback processing or financial exploitation vulnerability. Repeated assessments and residential histories would help address these questions. Our self-report measures capture perceived support, cognitive difficulties, and vulnerability rather than directly observed social resources, objective cognitive performance, or confirmed exploitation; future work could combine these measures with behavioral and longitudinal assessments. Finally, the neuroimaging analyses were exploratory, and cluster correction was applied within each statistical map rather than across the full set of models. Regional follow-ups describe the clusters identified in this sample and require independent evaluation. Replication in a larger, geographically diverse sample would help establish the generalizability of these associations.

Overall, the present study highlights the importance of considering environmental context and perceived social support when investigating social reward processing and financial exploitation vulnerability in older adulthood. Greater PM2.5 exposure was associated with higher financial exploitation vulnerability among individuals reporting lower perceived social support, while neighborhood deprivation and social support jointly related to neural responses to positive social and monetary feedback near the TPJ. Additionally, individual differences in financial exploitation vulnerability were associated with ventral striatal connectivity during social relative to monetary feedback, although this association requires further investigation. These findings suggest that environmental exposures may have distinct relationships with financial vulnerability and neural responses to social reward, and that considering perceived social support can reveal patterns that may be obscured when examining environmental factors independently. More broadly, understanding how environmental conditions and social resources intersect with the neural systems involved in processing social information may help clarify individual differences in vulnerability during older adulthood.

## Acknowledgments

This work was supported by the National Institute on Aging of the National Institutes of Health (R01-AG067011 to DVS). We thank Avi Dachs and Jamie-Nicole Luistro for assistance with data collection and Jimmy Wyngaarden for assistance with analyses.

